## Supplementary figures and images for "Computational Modeling of the Chlamydial Developmental Cycle Reveals a Potential Role for Asymmetric Division"

### Supplemental fig 1

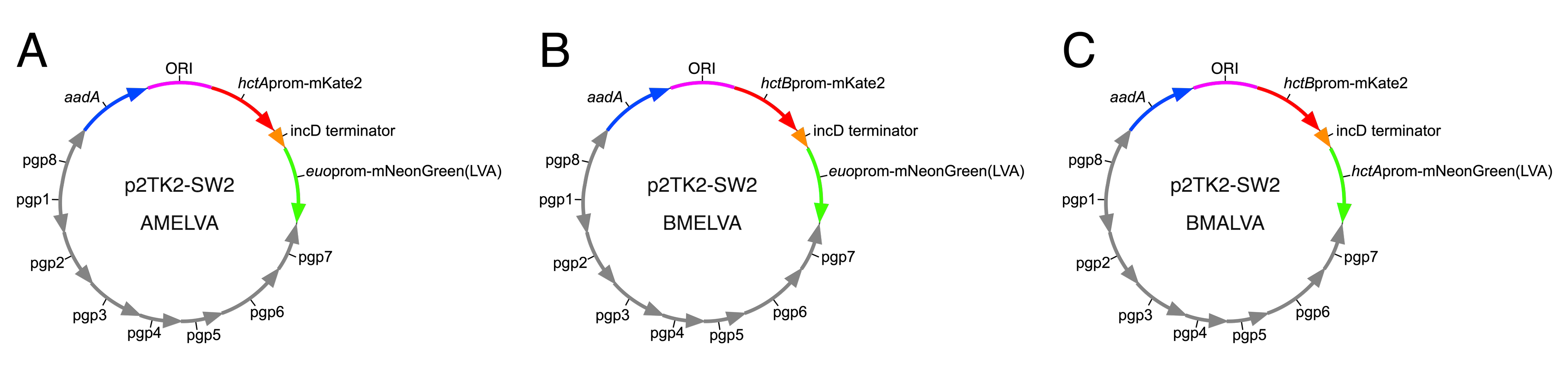

### Supplemental fig 2

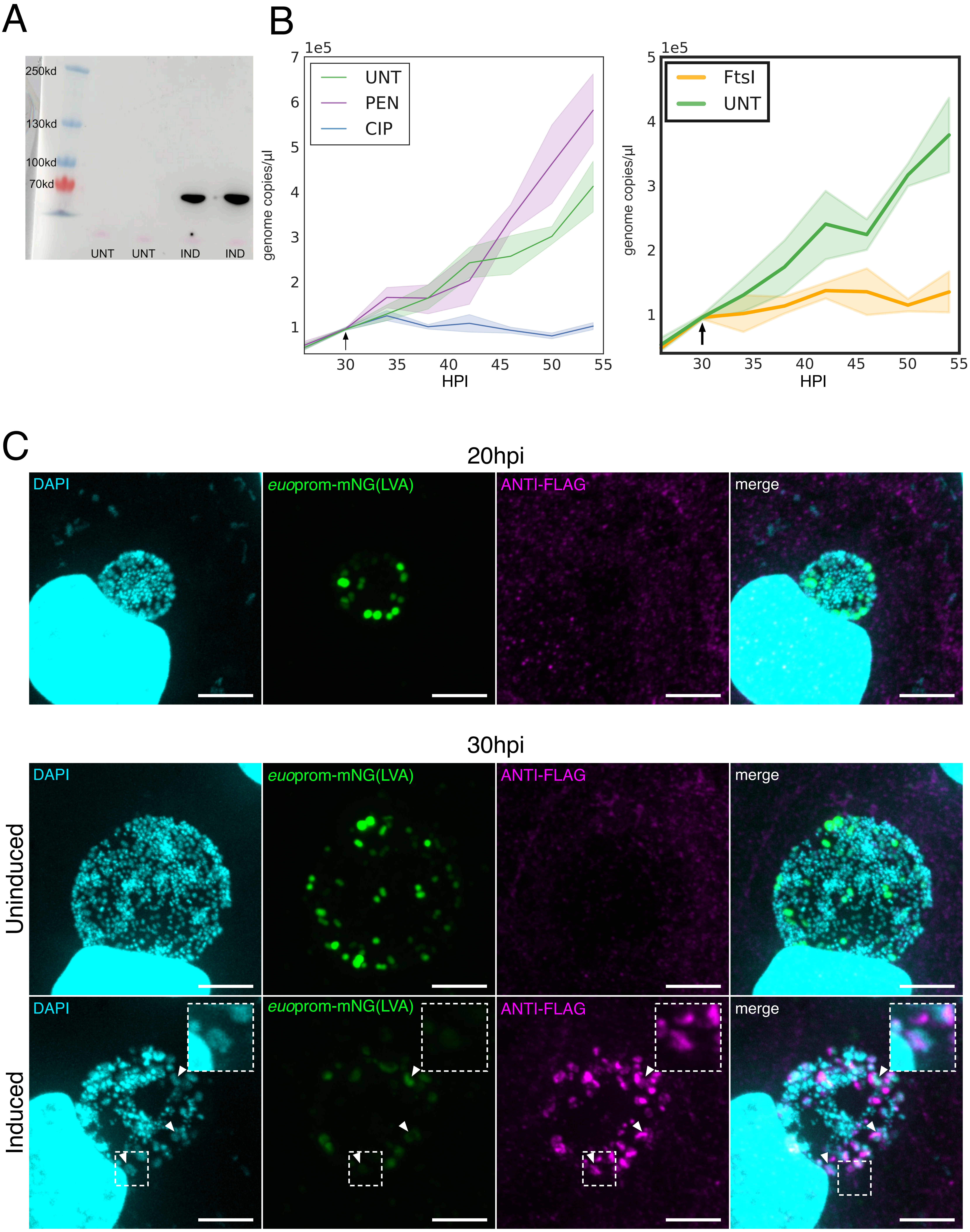

### Supplemental fig 3

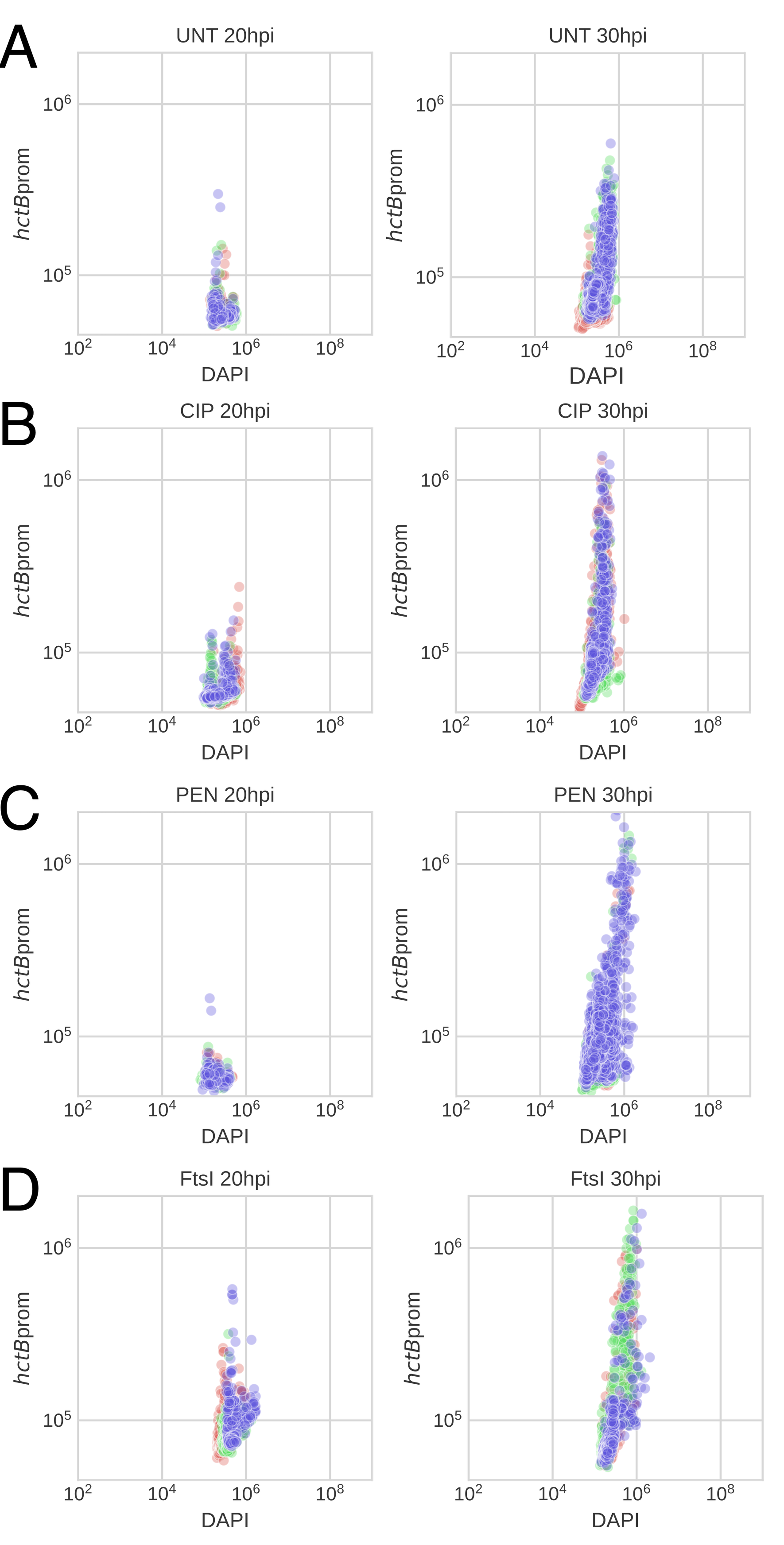

### Supplemental fig 4

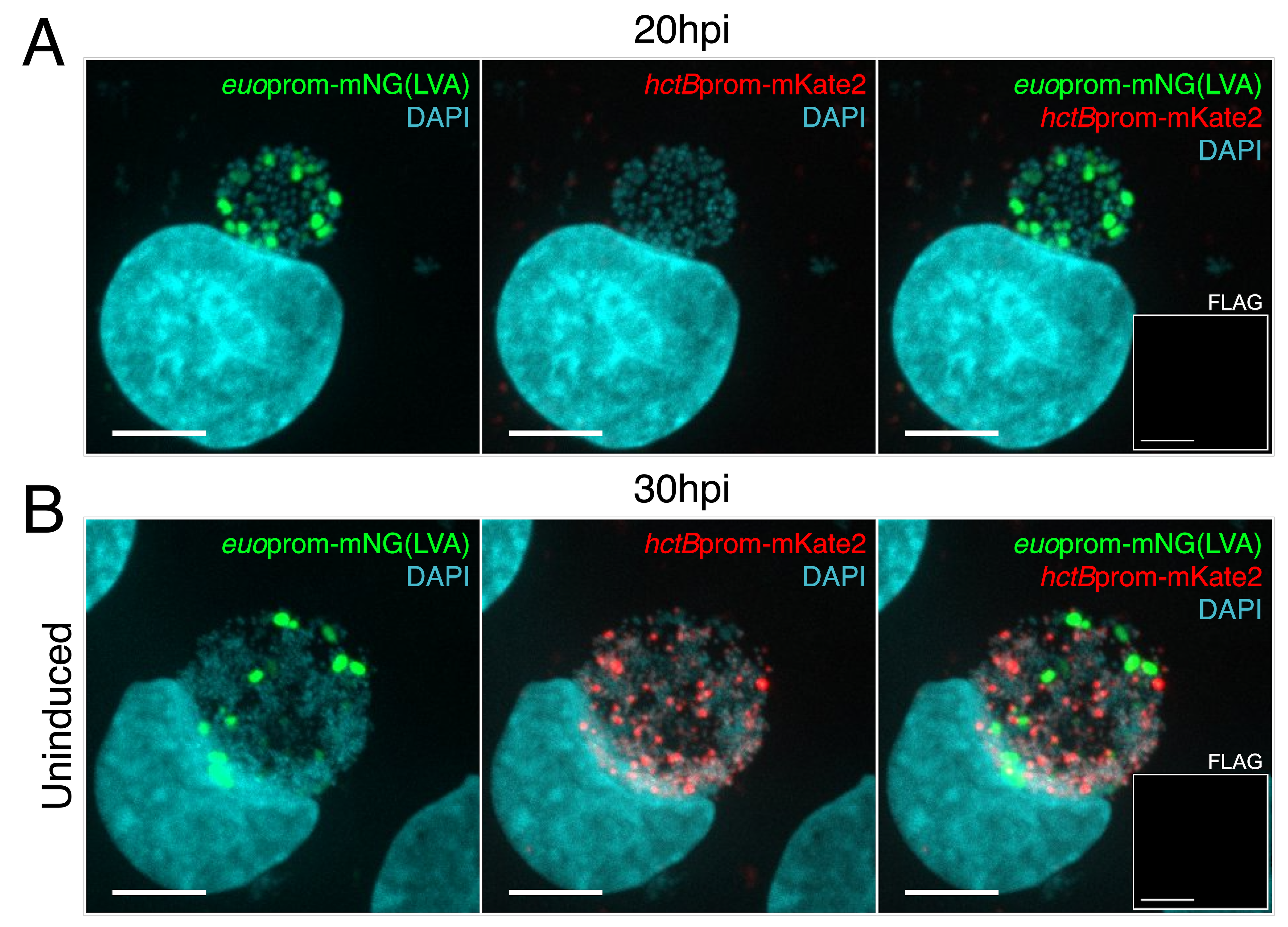
