## Supplemental model description for "Computational Modeling of the Chlamydial Developmental Cycle Reveals a Potential Role for Asymmetric Division"

**Supplemental Information: Description of Agent-based Models**

### Model Description

Two agent-based models that simulate the developmental dynamics of *Chlamydia* at the single inclusion level were created using the Python-based platform Cellmodeller. Agent-based models simulate and track individual cells (agents) within large complex population structures. Cellmodeller is a framework that models cellular biophysics, gene regulation and intercellular signaling (1). Models are created within this framework using simple model definitions with Python. Cellmodeller tracks the properties of each cell during the simulations. For our models, the physical information (position, size), growth rates, and expression levels of HctA, HctB and Euo were tracked.

EB germination, RB amplification, IB conversion/production, and EB formation were simulated for both models. A germination time of 10 ± 2 hours was used, and a doubling time (*R*) of RBs was estimated to be ~2 ±.05 hours based on our kinetic experiments. The two models differ in the mechanisms that generate the IB cell type.

### Creation of the IB

#### Direct Conversion model

In this model, RBs replicate symmetrically after germination to produce two RBs. Over time each new RB has an increasing chance of directly converting into an IB cell. We fit a sigmoidal function to the mean value of inclusion-level live-cell *euo*prom-mNG(LVA) expression data (Fig. 1). The function was set to scale between zero and one hundred percent and used to drive the percent chance of RB to IB conversion. Conversion was limited to a maximum of 50% chance of IB conversion as higher or lower values led to RB extinction or RB overproduction.

#### Asymmetric Production model

The asymmetric conversion model uses a maturation mechanism to switch from RB amplification to IB production. After germination, the initial RB ($RB_R$) divides symmetrically, producing two RBs after every division. The $RB_R$ matures into the $RB_E$ cell which acts as a stem cell producing an IB and regenerating the $RB_E$ after every division. For this model we used the same sigmoidal function to drive $RB_R$ to $RB_E$ maturation.

#### RB conversion function and parameterization

As the mechanism that drives either RB amplification/maturation or direct conversion is still unknown, we fit the following sigmoidal function (1) to the mean value of inclusion-level live-cell *euo*prom-mNG(LVA) expression data using the optimize.curve_fit function from the SciPy package. The parameters for this function were adjusted to scale between zero and one hundred percent (Supplemental Methods, Table 1) and used to drive the percent chance of $RB_R$ to $RB_E$ maturation (*M*) or chance of direct conversion for individual RBs (Supplemental Methods, Figure 1).

$$M = \frac{L}{1 + e^{(m-(tR))c} + d}$$

| Parameter | Value |
|---|---|
| *m* | 2.15841312E+01 |
| *L* | 97.81 |
| *c* | 6.77630536E-01 |
| *d* | 2.19 |

Supplemental Methods, Table 1: Parameters for RB maturation sigmoidal function.

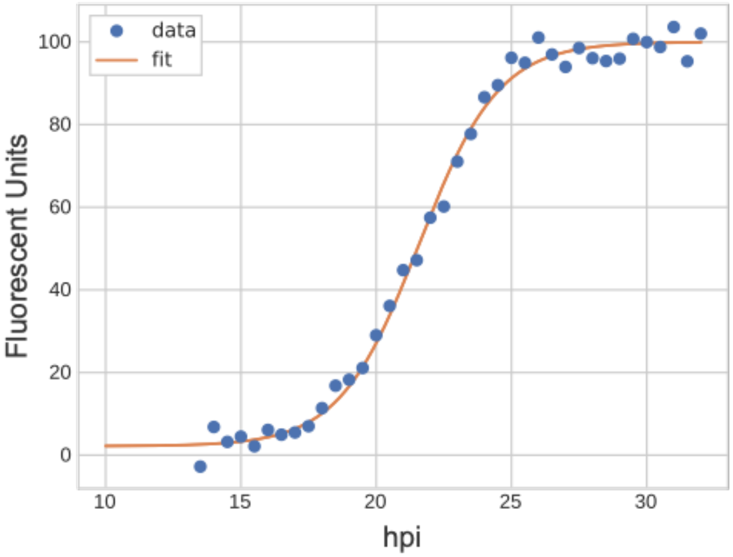


**Supplemental Methods, Figure 1: Sigmoidal function fit to live-cell *euo*prom-mNG(LVA) data.** Inclusion-level data was collected from host cells infected with L2-BMELVA. Fluorescent intensity was collected every 15 min using automated live-cell microscopy. A sigmoidal function (orange line) was fit to the mean of the live-cell data from 10 to 32 hpi and then scaled from zero to one hundred percent.

**Gene expression and EB formation**

The models also simulate RNA transcription and protein production dynamics of Euo ($E_R$, $E_P$), HctA ($A_R$, $A_P$), and HctB ($B_R$, $B_P$) for each cell form. The kinetics and cell specificity for the expression of each of these proteins was based on our live-cell inclusion-level and single-cell data (**Fig. 1 and 2, main text**). For both models, RBs express Euo but not HctA or HctB. After IB formation the Euo production rate is set to zero and when Euo protein levels drop below a threshold due to degradation, HctA expression is de-repressed. When HctA protein concentration reaches a specified threshold, HctB expression is induced. Once HctB levels reach a specified concentration threshold the cell is considered an EB and infectious. The following equations are used to drive RNA and protein expression in each cell form. The parameters for the equations are given in Supplemental Methods, Table 2.

$$E_R = E_R + (pr_E R) - (nr_E E_R R)$$

$$E_P = E_P + (p_E E_P R) - (n_E E_P R)$$

$$A_R = A_R + (pr_A R) - (nr_A A_R R)$$

$$A_P = A_P + (p_A A_P R) - (n_A A_P R)$$

$$B_R = B_R + (pr_B R) - (nr_B B_R R)$$

$$B_P = B_P + (p_B B_P R) - (n_B B_P R)$$

| **Parameter** | **Description** | **Value** |
|---|---|---|
| $pr_E$ | *euo* RNA production rate | 0.02 |
| $pr_A$ | *hctA* RNA production rate | 0.04 |
| $pr_B$ | *hctB* RNA production rate | 0.06 |
| $nr_E$ | *euo* RNA degradation rate | 0.02 |
| $nr_A$ | *hctA* RNA degradation rate | 0.01 |
| $nr_B$ | *hctB* RNA degradation rate | 0.024 |
| $p_E$ | *euo* protein production rate | 0.5 |
| $p_A$ | *hctA* protein production rate | 1.0 |
| $p_B$ | *hctB* protein production rate | 0.5 |
| $n_E$ | *euo* protein degradation rate | 0.08 |
| $n_A$ | *hctA* protein degradation rate | 0.05 |
| $n_B$ | *hctB* protein degradation rate | 0.01 |

Supplemental Methods, Table 2: Parameters used for expression dynamics.

**Modeling cell division inhibition**

To determine the effects of cell division inhibition on each cell type subpopulation, we simulated RB conversion to an alternative cell type where the RB was prevented in its ability to divide. For the direct conversion model, the RBs incapable of cell division were still permitted to convert directly into IBs based on the previously established rate of replication (conversion decision time equalling ~2 hours).

**Modeling RB cell death**

To determine the effects of RB cell death in both models the RBs were again classified as a separate cell type and removed from the RB population. However, this new cell type was inhibited in its ability of IB conversion/production regardless of the model.

The python model descriptions are available on GitHub at SGrasshopper/Chlamydial-developmental-cycle

**References:**

1. Rudge TJ, Steiner PJ, Phillips A, Haseloff J. Computational modeling of synthetic microbial biofilms. Vol. 1, ACS Synthetic Biology. American Chemical Society (ACS); 2012. p. 34552.
